# Elevated transcriptional pausing of RNA polymerase II underlies acquired resistance to radiotherapy

**DOI:** 10.1101/2021.10.11.462362

**Authors:** Honglu Liu, Chunhong Yu, Na Zhang, Yang Meng, Canhua Huang, Chunhong Hu, Fang Chen, Zhiqiang Xiao, Zhuohua Zhang, Hao Shao, Kai Yuan

**Author notes:** Corresponding author: Kai Yuan Ph.D., 110 Xiangya Road, Changsha, Hunan, China.

## Abstract

As the mainstay modality for many malignancies, particularly inoperable solid tumors such as nasopharyngeal carcinoma (NPC), ionizing radiation (IR) induces a variety of lesions in genomic DNA, evoking a multipronged DNA damage response to interrupt many cellular processes including transcription. The turbulence in transcription, depending on the nature of DNA lesions, encompasses local blockage of RNA polymerase II (RNAPII) near the damage sites, as well as a less understood genome-wide alteration. How the transcriptional change influences the effectiveness of radiotherapy remains unclear. Using a panel of NPC and lung cancer cell lines, we observe increased phosphorylation at serine 5 (pS5) of the RNAPII after IR, indicating an accumulation of paused RNAPII. Remarkably, a similar increase of pS5 is seen in IR-resistant cells. ChIP-seq analysis of RNAPII distribution confirms this increased pausing both in IR-treated and IR-resistant NPC cells, notably on genes involved in radiation response and cell cycle. Accordingly, many of these genes show downregulated transcripts abundance in IR-resistant cells, and individual knockdown some of them such as TP53 and NEK7 endows NPC cells with varying degrees of IR-resistance. Decreasing pS5 of RNAPII and hence tuning down transcriptional pausing by inhibiting CDK7 reverses IR-resistance both in cell culture and xenograft models. Our results therefore uncover an unexpected link between elevated transcriptional pausing and IR-resistance. Given the recurrent NPC tissues display a steady increase in pS5 compared to the paired primary tissues, we suggest that CDK7 inhibitors can be used in combination with radiotherapy to increase sensitivity and thwart resistance.

## Introduction

Radiotherapy is the mainstay treatment of non-metastatic solid tumors such as nasopharyngeal carcinoma, a geographically unbalanced malignancy prevalent mostly in southeast Asia including southern provinces of China (1). High-energy ionizing radiation (IR) causes tumor regression by generating a variety of lesions to living cells. In addition to oxidative damages, every Gray (Gy) of IR induces roughly 1000 single-strand breaks (SSBs) and 40 double-strand breaks (DSBs) to the nuclear DNA, eliciting most of the cytotoxicity after IR exposure (2,3). Comprehensive characterization of cellular responses to IR-induced DNA damage has been the central theme in the field to improve treatment efficacy and limit the development of resistance.

As the rate-limiting step of gene expression, the dynamic process of transcription underlies many cellular responses towards intrinsic and extrinsic stimuli. A full transcription cycle consists of four sequential phases—initiation, pausing, elongation, and termination—which are regulated by post-translational modifications on the C-terminal domain (CTD) of the Rpb1 subunit of RNA polymerase II (RNAPII) (4). After initial recruitment to promoter regions by transcription factors (TFs) and general transcription factors (GTFs), RNAPII CTD is phosphorylated at position 5 (pS5) of the Y_1_S_2_P_3_T_4_S_5_P_6_S_7_ repeat by the TFIIH-associated kinase CDK7, escaping from the promoter and then quickly becoming paused after transcribing 20-60 nucleotides downstream of the transcription start site (TSS) (5,6). This promoter-proximal pausing of RNAPII is a nexus of regulation, ensuring precise spatiotemporal control of gene activity in many biological processes, particularly in embryonic development (6–8). Several proteins have been identified to govern the release of the paused RNAPII, which includes the negative elongation factor (NELF) complex and the positive transcription elongation factor b (P-TEFb). The CDK9 subunit of P-TEFb along with other CDKs can phosphorylate NELF and the position 2 (pS2) of the CTD, thereby driving the paused RNAPII to enter the elongation phase (4,6,9). The RNAPII then continues traveling along the DNA template till it encounters the termination signal to complete the transcription cycle.

Transcription is highly sensitive to DNA lesions. Unscheduled encounters with damages on the DNA template can lead to transcriptional failure and genome instability. Therefore, cells deploy multiple ways to synergize transcription and DNA damage response (DDR) (9). RNAPII functions as a damage sensor during the repair of bulky single-strand DNA damages induced by ultraviolet light (UV). UV damage triggers an immediate release of paused RNAPII into gene bodies to detect local lesions and facilitate transcription-coupled nucleotide excision repair (TC-NER) (10). Following this initial phase, the RNAPII is then degraded by ubiquitination which leads to a global shutdown of transcription (9). DNA DSBs can also regulate transcription, inducing the formation of a repressive chromatin state mediated by histone H2AK119ub around the damaged site to inhibit local RNAPII activity (11–13). The global impact of DSBs on transcription dynamics however is less clear.

Radiotherapy induces both SSBs and DSBs in the target cells (3). How IR exposure modulates the global transcriptional dynamics and whether the altered kinetics in transcription cycle influences the treatment efficacy are outstanding questions await investigation. Here we show that irradiation leads to an increase in transcriptional pausing, and the elevated pausing level contributes to the development of IR-resistance.

## Materials and Methods

### Cell culture and drug treatment

The NPC cell lines (CNE2, Hone1, C666) and the lung cancer cell lines (H1264, H460) were obtained from Prof. Zhiqiang Xiao and Prof. Rong Tan (Xiangya Hospital, Central South University). The corresponding IR-resistant cell lines were prepared as previously described (14). NPC cell lines were cultured with DMEM/High Glucose medium (Biological Industries) containing 10% FBS (VISTECH, SE100-B), and lung cancer cell lines with RPMI 1640 medium (Biological Industries) containing 10% FBS, in a humidified incubator with 5% CO_2_ at 37 °C. Small molecular inhibitors THZ1 (Selleck, S7549), BS-181 (Selleck, S1572), KU-55933 (MedChemExpress, HY-12016), and PI-103 (Selleck, S1038) were used at 0.25 μM, 10 μM, 10 μM, and 0.4 μM respectively.

### Construction and transfection of shRNA plasmids

The shRNA oligos were synthesized by Tsingke (Tsingke Biotechnology) and cloned into pLKO.1-TRC vector. The sequences of shRNA were listed in Supplementary Table S1.

### Tumor cell viability analysis after irradiation

To confirm IR-resistance, cells were seeded into 96-well plates at the density of 1000-2000 per well. After 24 hours, cells were irradiated with the indicated X-ray dose at room temperature at 200 cGy/min with a linear accelerator (X-Rad 225, Precision). The viability was either measured daily using MTT (Sigma, M5655) for 6 days, or measured at day 5 after irradiation. For western blotting experiments, cells were harvested at day 2 after IR treatment.

For xenograft tumor model, 5×10^6^ of the indicated cells were suspended in 100 μL DPBS (Biological Industries) and injected subcutaneously into the flank of 6 weeks old female BALB/c nude mice (Hunan SJA Laboratory Animal). The tumors were measured thrice weekly with a digital caliper, and the volume calculated using the formula: 0.5 × (length × width^2^). Once the tumors reach 200 mm^3^ in size, the mice were irradiated with a single dose of 4 Gy under 2,2,2-tribromethanol anesthesia (Avertin). Only the tumor was irradiated, and the rest of the body was shielded by lead. All the animal experiments were approved by the Medical Ethics Committee of Central South University, and conducted according to the Guidelines of Animal Handling and Care in Medical Research in Hunan Province, China.

### Western blotting

Cells were washed with cold DPBS (Biological Industries) two times and then lysed in sample buffer (2% SDS, 10% glycerol, and 62.5 mM Tris-HCl, pH 6.8) supplemented with 1× protease inhibitor cocktail (Sigma, P8340), sodium fluoride (10 mM, Sigma, 450022), and sodium orthovanadate (1 mM, Sigma, 450243). After sonication, the protein concentration was determined using BCA assay (Beyotime, P0009). The following primary antibodies were used: mouse anti-RNAPII Ab (1:3000, Abcam, ab817), mouse anti-RNAPII phosphor S5 Ab (1:3000, Abcam, ab5408), rabbit anti-RNAPII phosphor S2 Ab (1:5000, Abcam, ab5095), mouse antitubulin HRP Conjugate (1:5000, Cell Signaling Technology, 12351S). The corresponding secondary antibodies (Thermo Fisher Scientific) were used at 1:5000. The signal was detected with ECL substrates (Millipore, WBKLS0500).

### ChIP-seq and ChIP-qPCR

The cells in a 100 mm dish were cross-linked with 1% formaldehyde for 10 min, quenched with 125 mM glycine for 5 min, rinsed with ice-cold PBS twice, and scraped in 1 mL PBS. After spinning at 1350 g for 5 min at 4□, the pellet was resuspended in 500 μL Lysis Buffer I (50 mM HEPES-KOH, pH 7.5, 140 mM NaCl, 1 mM EDTA, 10% glycerol, 0.5% NP-40, 0.25% Triton X-100), and incubated at 4□ for 10 min with rotating. After spinning at 4□, the pellet was then resuspended in 500 μL Lysis Buffer II (10 mM Tris-HCl, pH 8.0, 200 mM NaCl, 1 mM EDTA, 0.5 mM EGTA), and incubated for 10 min at room temperature. After spinning at 4□, the pellet was resuspended in 500 μL Lysis Buffer III (10 mM Tris-HCl pH 8.0, 100 mM NaCl, 1 mM EDTA, 0.5 mM EGTA, 0.1% Na-Deoxycholate, 0.5% N-lauroylsarcosine). The cell lysate was sonicated till the DNA was sheared to 200 bp. The lysate was then quenched by 1% of Triton X100, supplemented with 1 × protease inhibitor, and spun at 12000 rpm for 10 min at 4□. 50 μL supernatant was reserved as input, and the rest was incubated overnight at 4□ with the magnetic beads bound with 4 μg anti-RNAPII (Abcam, ab817), anti-RNAPII phospho S5 (Abcam, ab5408), or anti-RNAPII phospho S2 (Abcam, ab5095). The beads were washed four times with Wash Buffer (50 mM HEPES-KOH, pH 7.6, 500 mM LiCl, 1 mM EDTA, 1% NP-40, 0.7% Na-deoxycholate). The DNA was eluted with 100 μL of Elution Buffer (50 mM Tris-HCl, pH 8.0, 10 mM EDTA, 1% SDS). The cross-links were reversed by incubating in a heating oscillator at 65□ for 2 hours. The sample was incubated with 4 μL of 25 mg/mL RNase A overnight at 65□, and then with 2 μL of 10 mg/mL proteinase K at 55°C for 4 hours. The DNA was purified by phenol:chloroform:isoamyl alcohol extraction and resuspended in 20 μL ddH_2_O. ChIP-seq library was prepared using NEBNext® ChIP-Seq Library Prep Reagent Set (New England Biolab, E7645S). The library was sequenced on Illumina NovaSeq 6000 (Novogene). ChIP-qPCR was performed using the SYBR Green qPCR Master Mix (SolomonBio, QST-100) on the QuantStudio 3 Real-Time PCR system (Applied Biosystems). Primers were listed in Supplementary Table S1.

ChIP-seq raw reads were filtered using trim_galore v0.6.0, aligned to the hg38 genome assembly using Bowtie2 v2.3.5.1 with default parameters. Duplicate reads were removed using MarkDuplicates from the gatk package v.4.1.4.1. The pausing index (PI) was calculated as previously described (15). The RefSeq gene model was downloaded from UCSC. ChIP-seq and input reads were calculated using bedtools coverage v2.26.0, mapped to the TSS regions (TSSR, −50 to +300 bp relative to TSS) and the gene bodies (TSS +300 bp to +3 kb past the TES) for each annotated RefSeq isoform. The reads density was normalized by the region length and by the mapped filtered read numbers multiplied by 1 million (rpm/bp). The input was then subtracted and PI was calculated as the ratio between RNAPII density in the TSSR and the gene body. For multiple RefSeq isoforms of the same gene, the one with the strongest RNAPII ChIP-seq signal in the TSSR, at least 0.001 rpm/bp, was selected. All those genes with PI >2 were defined as paused, and the rest were non-paused. The bigwig tracks were generated using bamCompare from deeptools. Negative values were set to zero. IGV v.2.4.13 was used to visualize the bigwig tracks. ChIP-seq profiles were created by computeMatrix and plotProfile in deeptools.

### RNA-seq and RT-qPCR

RNA was extracted using TRIzol (Life Technologies, 87804). Libraries were prepared using mRNA-Seq Sample Preparation Kit (Illumina) and sequenced on an Illumina NovaSeq platform (Novogene). Raw reads were filtered using trim_galore, then mapped to hg38 genome assembly using STAR v2.7.1a. Differential expression analysis was performed using the Bioconductor package DESeq2. For RT-qPCR, RNA was converted to cDNA using the PrimeScript RT reagent Kit (Takara, RR037A).

### Immunofluorescence and live cell imaging

Cells were quickly rinsed with pre-warmed DPBS, fixed with pre-cooling methanol for 10 min at −20°C, and permeabilized in 0.5% Triton X-100 for 10 min. Cells were then blocked with 5% Bovine Serum Albumin for 30 min and incubated overnight at 4°C with anti-RNAPII phospho S5 (1:1500, Abcam, ab5408), anti-RNAPII phospho S2 (1:3000, Abcam, ab5095), anti-γ-Tubulin (1:1000, Sigma, T6557), anti-α-Tubulin (1:1000, Abcam, ab18251), or anti-ACA human centromere antibody (1:3000, gifted by Prof. Xuebiao Yao at University of Science and Technology of China). Cells were rinsed with DPBS thrice and incubated at room temperature for 1 hour with Alexa Fluor 488, 546, or 647 secondary antibodies (1:400, Thermo Fisher Scientific). For frozen sections, mouse tumor tissue was immediately embedded in optimum cutting temperature media (SAKURA Tissue-Tek® O.C.T. Compound 4583) and frozen at −80°C. Cryostat sections (12μm) were made and kept at −80°C. Slides were fixed in acetone for 5 minutes at room temperature and then rinsed with DPBS twice before being blocked with 5% BSA for 30 minutes. The slides were then incubated overnight at 4°C with anti-RNAPII phospho S5 (1:1000), rinsed thrice, and incubated for 1 hour at room temperature with Alexa Fluor 488 secondary antibody (1:400). DNA was visualized by DAPI staining (0.5 μg/mL, Sigma, D9542). The samples were mounted with SlowFade™ Diamond Antifade Mountant (Thermo Fisher Scientific, S36963).

For live imaging, cells in glass bottom dish (NEST, 801001) were transfected with pEGFP-Tubulin and mCherry-H2B using Lipofectamine 3000 (Invitrogen), and synchronized with double thymidine block procedure. Cells were then released into thymidine-free medium for 8 hours and imaged in PeCon environmental microscope incubator (ZEISS) at 37°C and 5% CO_2_. Images were collected on the LSM 880 confocal microscope using a 63× oil immersion objective lens with Airyscan mode (ZEISS).

### Immunohistochemistry (IHC)

All the human tissue related experiments were approved by the Medical Ethics Committee of Central South University, and the informed consent was obtained from the patients. IHC was performed to detect the level of RNAPII pS5 in5 pairs of nasopharynx cancer samples using RNAPII phosphor S5 antibody (1:10000, Abcam, ab5408). The IHC score was evaluated by two independent investigators blinded to the histopathologic features and clinical characteristics using the intensity and proportion of positively stained tumor cells as previously described (16).

### Statistical analyses

The experiments were carried out at least three times. Data were presented as the mean ± standard deviation (SD). Statistical analysis and survival fraction analysis was performed using GraphPad Prism 9. Flow cytometry data were analyzed using ModFit LT 4.1. The statistical details of each experiment were indicated in the respective figure legends. The two-tailed unpaired Student’s t-test or chi-squared test was performed to evaluate significant differences between the two groups. Kolmogorov-Smirnov test was used to compare significant differences between two groups of pausing index. P values are presented as star marks in figures: *p < 0.05, **p < 0.01, ***p < 0.001.

### Data availability

ChIP-seq and RNA-seq data will be submitted to the NCBI GEO database after acceptance. All custom scripts are available from the authors upon request.

## Results

### Increased RNAPII serine 5 phosphorylation both in IR-treated and IR-resistant cells

The paused RNAPII is enriched in pS5 phosphorylation and the elongating RNAPII pS2 (Fig. 1A). To compare the global transcriptional kinetics in radio-sensitive and -resistant cells, we assessed the phosphorylation status of RNAPII in NPC cell line CNE2 and its derivative radiation resistant cell line CNE2-IR (Fig. S1A). Results from western blot and immunofluorescence showed a consistent increase of pS5 as well as a slight decrease of pS2 in CNE2-IR cells compared with its parental CNE2 cells (Fig. 1B-1C). We expanded the analysis to two other NPC cell lines, two lung cancer cell lines, and their derivative radio-resistant cells (Fig. S1B). Except one lung cancer cell line H460, the radio-resistant cells all showed different levels of increase in pS5 RNAPII (Fig. 1D).

**Figure 1.**
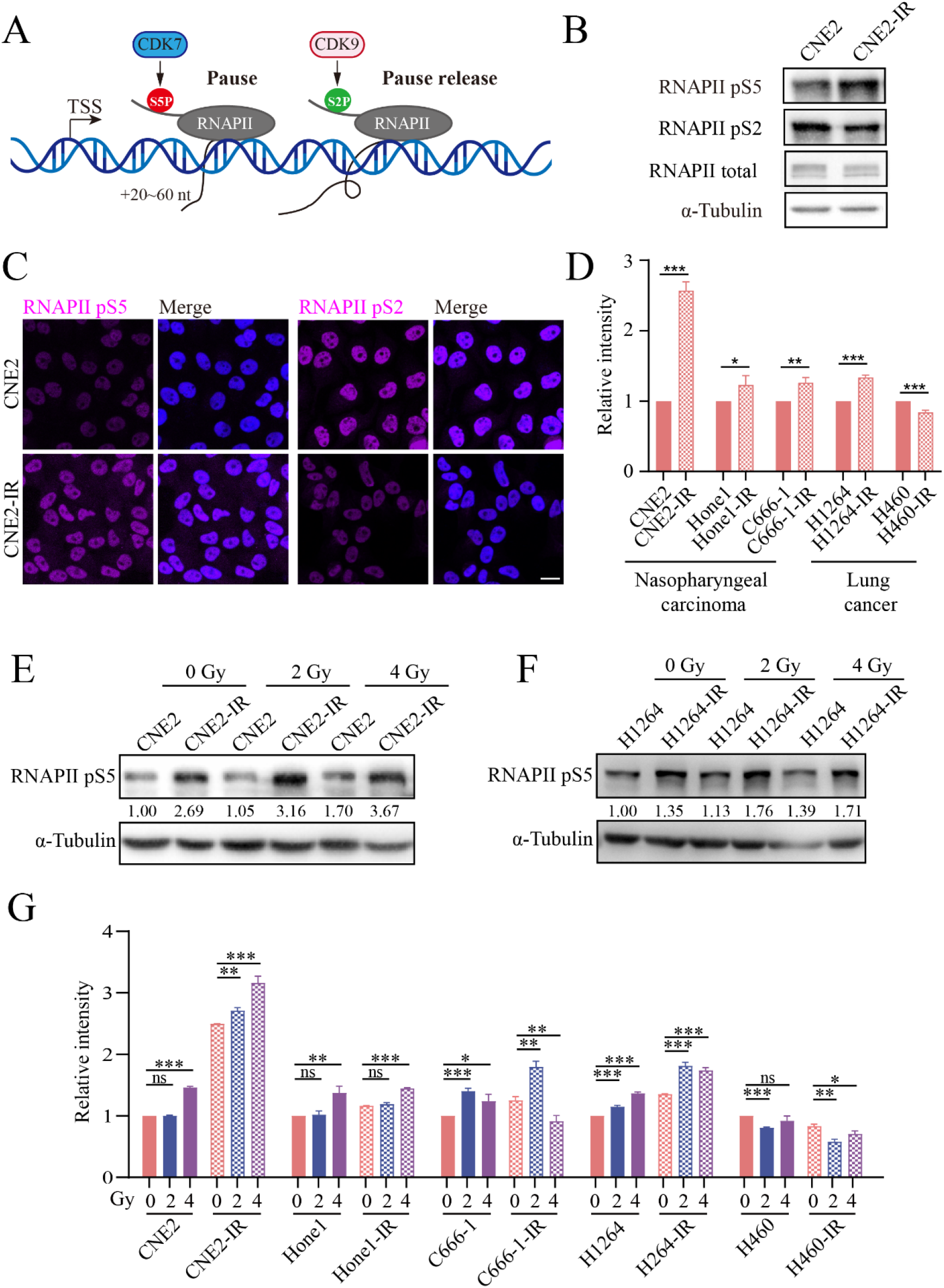
The RNAPII pS5 is increased both in IR-treated and IR-resistant cells. (A) Schematic of RNAPII CTD phosphorylation in transcription. (B) Western blot detecting RNAPII pS5 and pS2 levels in CNE2 and CNE2-IR cells, with α-tubulin as the loading control. (C) Immunofluorescence (purple) detecting RNAPII pS5 and pS2 in CNE2 and CNE2-IR cells. DNA is visualized by DAPI staining (blue). Scale bar:□20 μm. (D) RNAPII pS5 level in several nasopharyngeal and lung cell lines. Error bars indicate SD. *p < 0.05, **p < 0.01, ***p < 0.001 by t-test. (E-F) Western blot detecting RNAPII pS5 level in CNE2, CNE2-IR, H1264, and H1264-IR cell lines after receiving the indicated doses of irradiation. (G) Quantification of the RNAPII pS5 level in the indicated cells after radiation. Error bars indicate SD. *p < 0.05, **p < 0.01, ***p < 0.001 by t-test.

Radio-resistant cells are derived from parental cell lines by repeated exposure to IR. To test if IR treatment could increase the amount of pS5 RNAPII, we irradiated the cells with different radiation doses and detected the serine 5 phosphorylation. Most of the radio-sensitive and radio-resistant cell lines analyzed displayed increased pS5 after irradiation (Fig. 1E-1G, S1C-S1D). The H460 cells again showed a different response for reasons unknown (Fig. S1E).

These results showed that irradiation in many cells could elicit an increase of pS5 in the CTD of RNAPII, and this increase seemed to be persistent in the cells acquired IR-resistance, indicating that the kinetics of transcription was sensitive to IR and the pool of paused RNAPII was increased in IR-treated as well as IR-resistant cells. The DSBs caused by IR are central in IR-induced cellular effects (3). We inhibited the core kinases mediating the DSBs repair, ATM and DNA-PK (17), and no longer observed the increase of pS5 RNAPII after irradiation (Fig. S1F–S1G), suggesting that signaling pathways triggered by DSBs were responsible for the transcriptional change.

**Figure S1.**
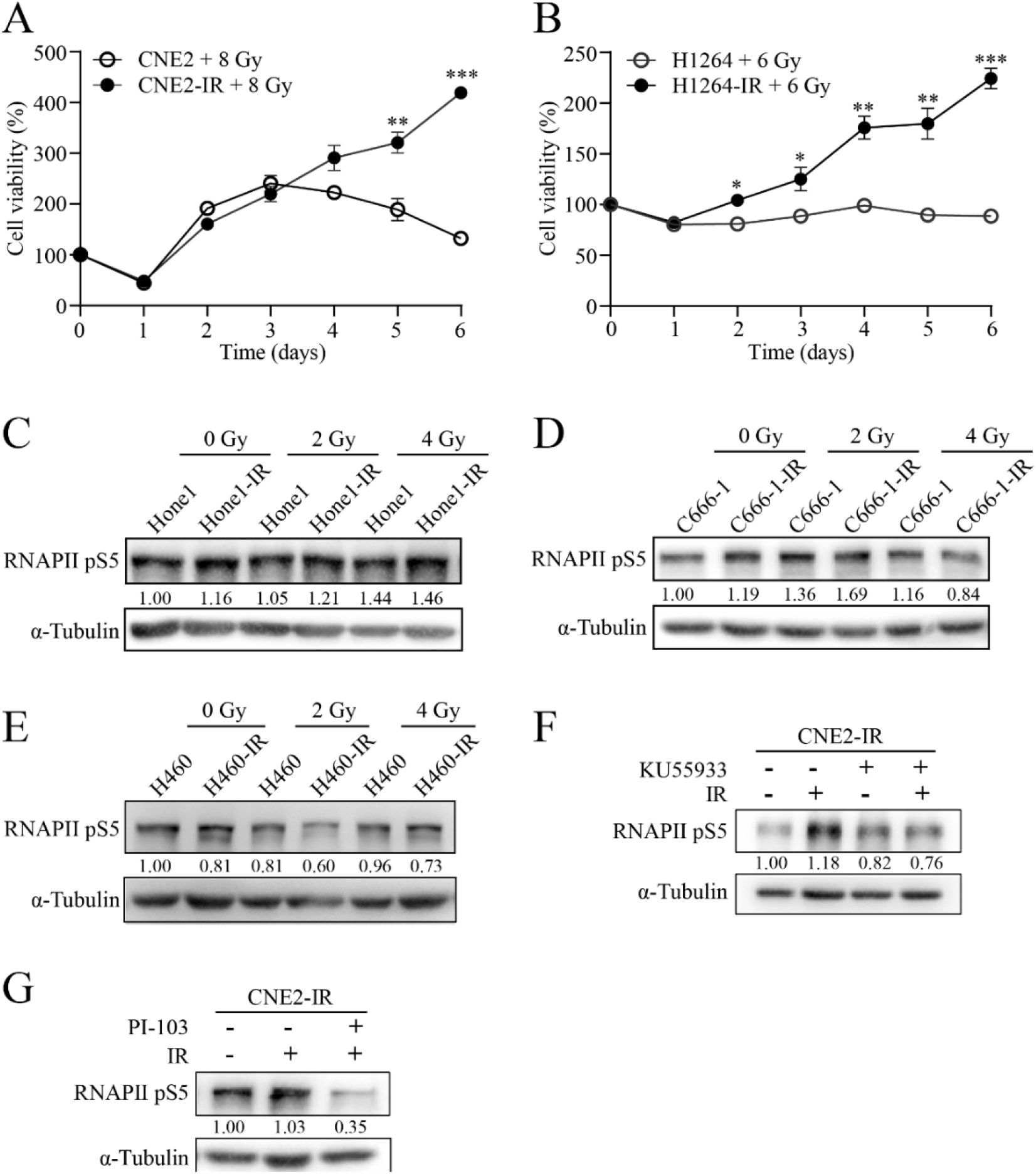
Measurement of RNAPII pS5 after radiation. (A-B) Measurement of cell viability after radiation using MTT assay. Data represent mean ± SD. *p < 0.05, **p < 0.01, ***p < 0.001 by t-test. (C-E) Western blot detecting RNAPII pS5 level in the indicated cell lines after different doses of irradiation. (F-G) Western blot detecting RNAPII pS5 level in CNE2-IR cells treated with radiation (4 Gy) in the presence of the indicated inhibitors.

### Elevated transcriptional pausing is associated with IR-resistance

To further elaborate the altered distribution of the transcribing RNAPII, we performed ChIP-seq analysis with CNE2, CNE2 cells after 4 Gy irradiation, as well as CNE2-IR cells (Fig 2A). Consistent with western blot and immunofluorescence results, we detected a global increase of pS5 around the TSS both in the irradiated CNE2 cells and the radio-resistant CNE2-IR cells even without irradiation (Fig. S2A–S2B). The pS2 of RNAPII was accordingly decreased at the TSS as well as in the gene bodies (Fig. S2C–S2D).

**Figure 2.**
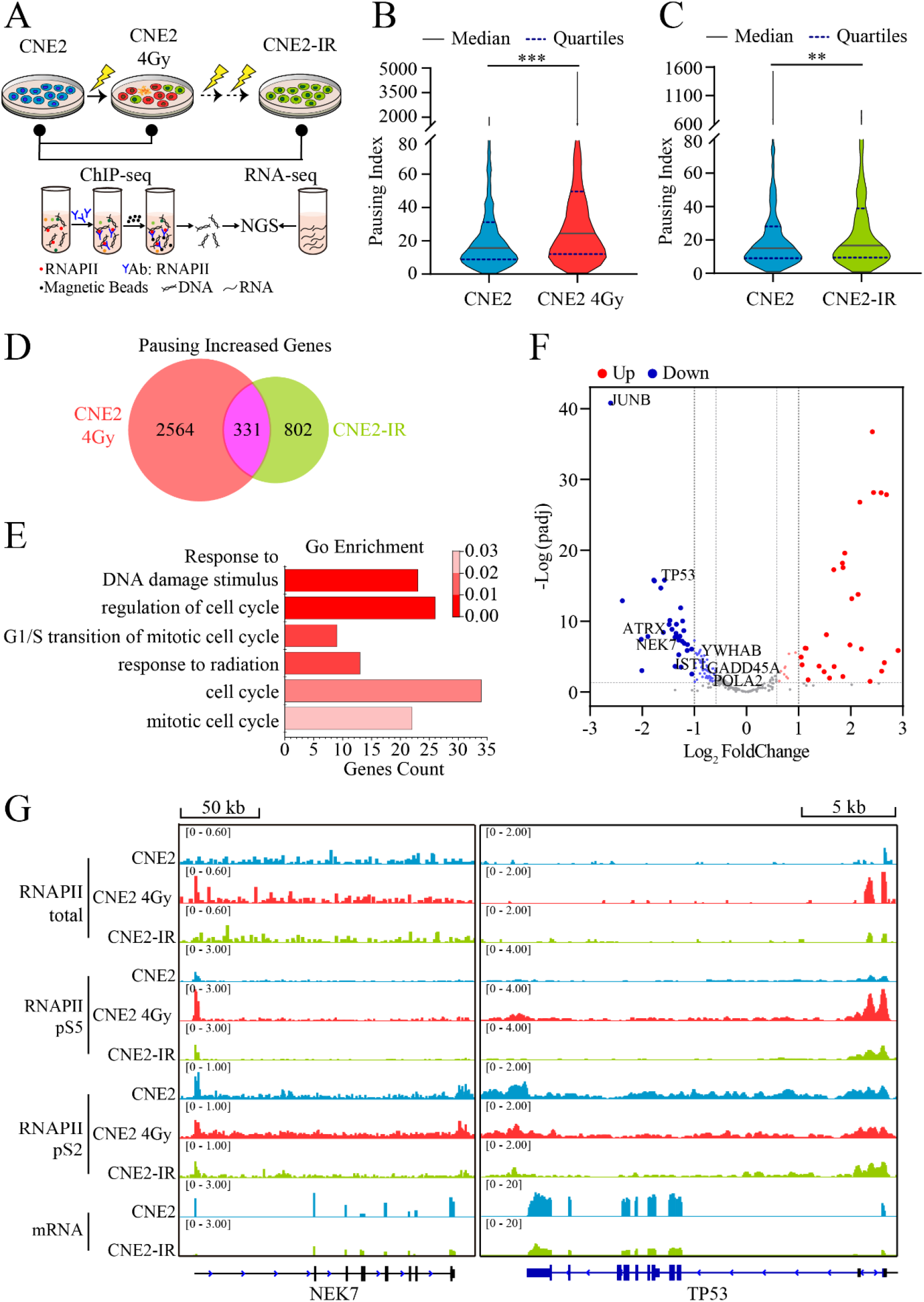
RNAPII pauses on many cell cycle genes in the radio-resistant cells. (A) Schematic showing the experimental designs for ChIP-seq and RNA-seq analyses. (B-C) Violin plots comparing RNAPII PI between CNE2 and CNE2 treated with 4 Gy irradiation, and CNE2 versus CNE2-IR cells. **p < 0.01, ***p < 0.001 by Kolmogorov-Smirnov test. (D) Venn diagram of RNAPII pausing increased genes. (E) Gene ontology (GO) analysis of the overlapping 331 genes. (F) Volcano plot showing the expression of the 331 pausing increased genes. Up (red) and down (green) regulated genes are determined with the cut-off values of adjusted p < 0.05 and |Log_2_FoldChange| > 0.585. (G) Genomic snapshots of ChIP-seq and RNA-seq signals at NEK7 and TP53 loci.

We further calculated the pausing index (PI) for each gene using ChIP-seq data generated with the total RNAPII antibody (Fig. S2E) (15). The overall PI in CNE2 cells received 4 Gy irradiation was significantly higher than that in control cells (Fig. 2B), and a similar increase in PI was observed in CNE2-IR cells (Fig. 2C and Supplementary Table S2). We identified 2895 genes in the irradiated CNE2 cells and 1133 genes in CNE2-IR cells that had increased PI compared with control CNE2 cells, with a total of 331 genes shared by these two lists (Fig. 2D and Supplementary Table S3). Gene ontology (GO) analysis of the 331 genes showed that these genes were enriched in biological processes including response to DNA damage, response to radiation, and cell cycle (Fig. 2E and Supplementary Table S4).

Change in transcriptional dynamics could affect the gene expression level. We performed RNA-seq analysis on CNE2 and CNE2-IR cells, and identified the differentially expressed genes. Of particular interest, in the 331 PI increased genes, 83 showed reduced expression in CNE2-IR cells (Fig. 2F and Supplementary Table S5). Many of these genes were involved in cell cycle regulation, such as TP53, NEK7, JUNB, etc. We selected the TP53 and NEK7 loci, designed primer sets targeting TSS and gene body regions according to the sequencing results (Fig. 2G), and validated by ChIP-qPCR that the pausing of RNAPII was indeed elevated on these genes (Fig. S2F–S2G).

Based on these results, we concluded that a significant fraction of RNAPII underwent increased promoter-proximal pausing after irradiation, and this increased pausing was maintained on a subset of genes involved in response to radiation as well as cell cycle, downregulating their mRNA levels in cells acquired radio-resistance.

**Figure S2.**
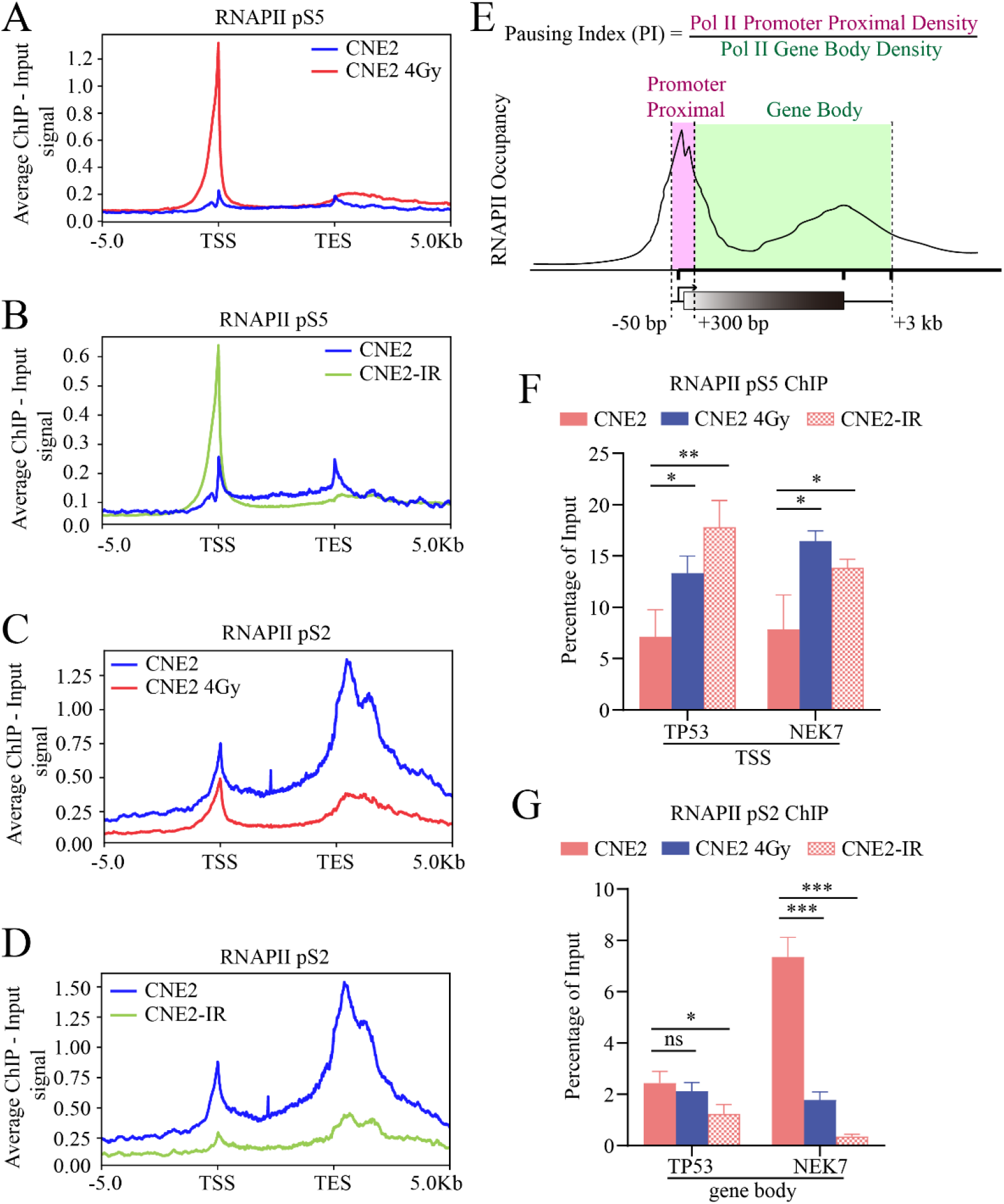
IR-induced RNAPII pausing. (A-B) Profiles of the pS5 RNAPII ChIP-seq signal on genomic regions spanning 5 kb upstream of the TSS and 5 kb downstream of the transcription end site (TES) of the human refseq genes in the indicated cells. (C-D) Profiles of the pS2 RNAPII ChIP-seq signal. (E) Calculation of PI using RNAPII ChIP-seq data (15). (F) ChIP-qPCR of RNAPII pS5 around the TSS of TP53 and NEK7 genes. (G) ChIP-qPCR of RNAPII pS2 in the gene body regions of the two genes. Data represent mean ± SD. *p < 0.05, **p < 0.01, ***p < 0.001 by t-test.

### Many pausing regulated genes are contributing to IR-resistance

To assess the contribution of these downregulated genes to the development of resistance to radiation, we individually knocked down the expression of several of them including TP53, NEK7, JUNB, PKN2, and POLA2 in the radio-sensitive CNE2 cells using small hairpin RNA (shRNA) (Fig. S3A). While the control CNE2 cells demonstrated a severe decline in viability in response to the increasing irradiation dose, the CNE2 cells treated with shRNA targeting different candidate genes all gained varying degrees of resistance to irradiation, with TP53 and NEK7 knockdown (KD) showing the highest resistance close to that of the CNE2-IR cells (Fig. 3A). Flow cytometry analysis revealed that the CNE2-IR cells had a slight G2/M arrest compared to CNE2 cells, and NEK7 KD in CNE2 cells caused a similar effect on the cell cycle (Fig. 3B).

**Figure 3.**
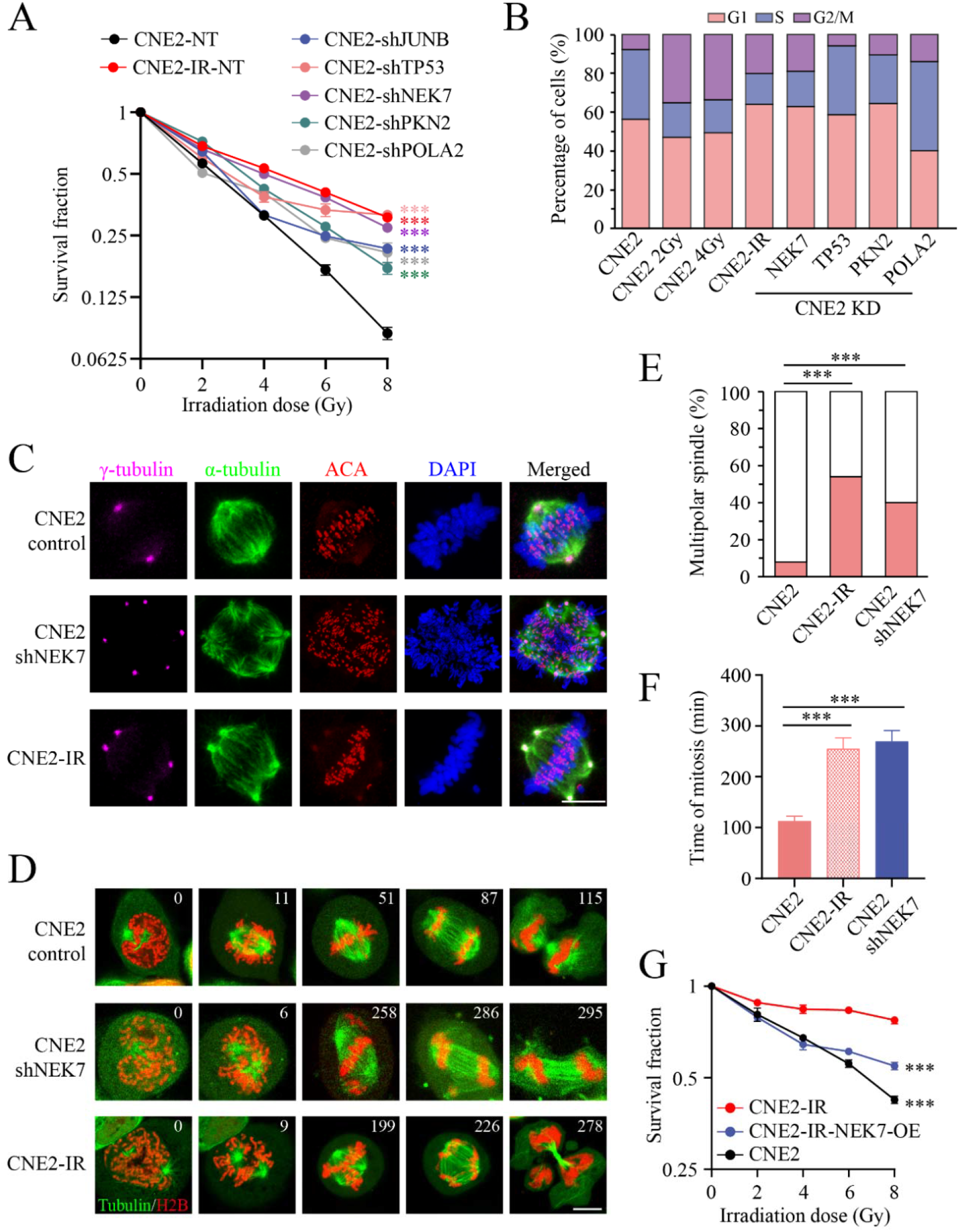
NEK7 as a new resistance factor regulated by IR-induced transcription pausing. (A) Cell survival curves of CNE2-IR and CNE2 cells infected with lentiviruses expressing control shRNA or shRNA targeting the indicated genes after irradiation with different doses. ***p < 0.001 by t-test. (B) Flow cytometry analysis of the cell cycle using propidium iodide. (C) Fluorescent staining of double-thymidine synchronized cells. Mitotic centrosomes are labeled with anti-γ-tubulin (purple), spindles are labeled with anti-α-tubulin (green), centromeres are labeled with anti-ACA (red), and chromosomes are labeled with DAPI (blue). Scale bar: 10 μm. (D) Confocal images from live cell imaging experiments. Chromosomes are visualized with mCherry-Histone 2B (red), and spindles with GFP-tubulin (green). Numbers on the top-right represent elapsed time. Scale bar: 10 μm. (E) Quantification of multipolar spindles. n = 50. ***p□<□0.001 by chi-squared test. (F) Quantification of the mitotic duration. Data represent mean ± SD (n > 4). ***p□<□0.001 by t-test. (G) Survival curves of CNE2, CNE2-IR, and CNE2-IR cells overexpressing NEK7 after irradiation with different doses. ***p < 0.001 by t-test.

We further characterized the cell cycle changes seen in the CNE2-IR cells and CNE2 cells treated with NEK7 shRNA. Immunofluorescence showed that both CNE2-IR and the NEK7 KD CNE2 cells displayed defects in the formation of bipolar mitotic spindle (Fig. 3C and S3C), with a ratio of multipolar spindle greatly exceeding that seen in the control CNE2 cells (Fig. 3E, 54% in CNE2-IR, 40% in CNE2 NEK7 KD, versus 8% in CNE2). To evaluate the influence of this spindle defect on the progression of mitosis, we performed live-cell imaging in the indicated cells using ectopically expressed GFP-tubulin and mCherry-Histone 2B to visualize mitotic spindle and chromosomes respectively (Fig. 3D). The duration of mitosis was about 2 hours in CNE2 control cells, whereas in CNE2-IR or NEK7 KD CNE2 cells, abnormal spindle formation was often observed and the mitotic phase was significantly extended (Fig. 3F, 113 ± 8 minutes in CNE2 cells versus 255 ± 18 minutes in CNE2-IR and 270 ± 19 minutes in CNE-2 NEK7 KD cells). In the most extreme cases, the cells were arrested in prometaphase for more than 300 minutes and could not finish mitosis (Fig. S3D).

The expression of NEK7 was reduced in the CNE2-IR cells. To test if the downregulation of NEK7 was important for maintaining the IR-resistance, we ectopically overexpressed NEK7 in the CNE2-IR cells (Fig. S3B), and we found that increase the expression of NEK7 attenuated the viability of the radio-resistant CNE2-IR cells after irradiation (Fig. 3G).

These results indicated that many of the genes regulated by IR-induced transcriptional pausing were contributing to the development of radio-resistance, and identified NEK7 as a new IR-resistance factor, likely via affecting mitotic centrosomes as previously reported (18,19).

**Figure S3.**
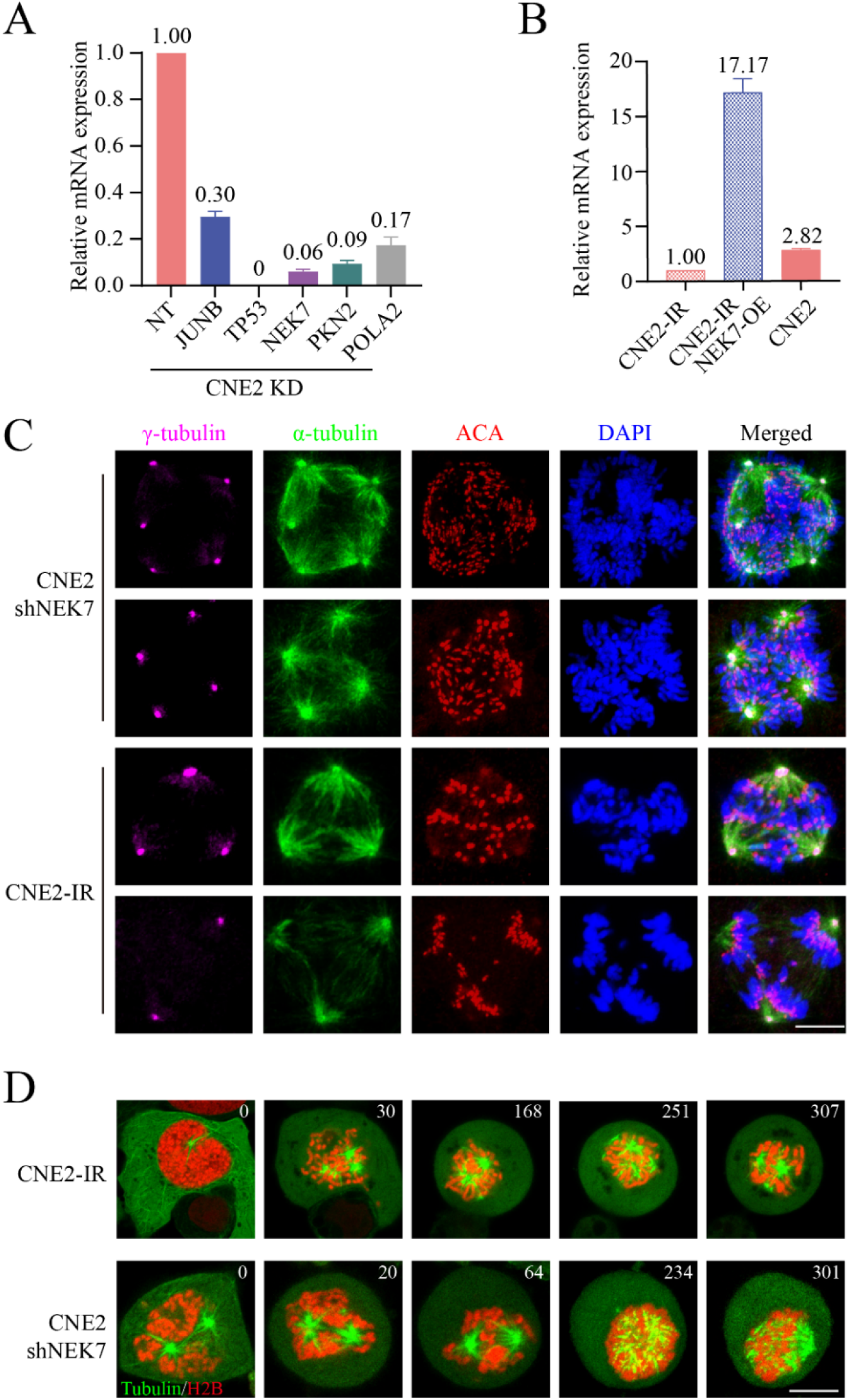
NEK7 knockdown results in multipolar spindle and mitotic arrest. (A) qPCR results showing the knockdown efficiency of different shRNA targeting the indicated genes. (B) qPCR results showing the mRNA levels of NEK7. Data represent mean ± SD. (C) Mitotic cells with multipolar spindles visualized by immunofluorescence. Mitotic centrosomes are labeled with anti-γ-tubulin (purple), spindles are labeled with anti-α-tubulin (green), centromeres are labeled with anti-ACA (red), and chromosomes are labeled with DAPI (blue). Scale bar: 10 μm. (D) Confocal images from live cell imaging experiments. Chromosomes are visualized with mCherry-Histone 2B (red), and spindles with GFP-tubulin (green). Numbers on the top-right represent elapsed time. Scale bar: 10 μm.

### Inhibiting transcriptional pausing reverses IR-resistance

Since many of the pausing regulated genes were able to modulate IR-resistance, we reasoned that targeting transcriptional pausing could be an effective strategy to restrain resistance to radiotherapy. CDK7 in the TFIIH is required for RNAPII pausing and CDK9 in pTEFb promotes transcriptional elongation. We first tested if CDK7 inhibition could influence sensitivity to radiation. We treated the CNE2 and CNE2-IR cells with two different CDK7 inhibitors, THZ1 and BS-181 (20,21). The CNE2-IR cells were more sensitive to THZ1 compared to CNE2 cells, with IC50 of 209.6 nM versus 1027.0 nM in CNE2 cells (Fig. 4A). We observed a reduced but consistent effect with BS-181 as well (Fig. S4A). We combined the chemical inhibition with radiation, and found that the joint treatment reversed the ratio-resistance of the CNE2-IR cells (Fig. 4B and S4B). The treatment with THZ1 or BS-181 also inhibited the increase of RNAPII pS5 induced by radiation (Fig. S4C and S4D). We next investigated if enhancing the activity of CDK9 could reduce resistance. Hexim1 is a key component of the ribonucleoprotein (RNP) complex inhibiting pTEFb (22). We knocked down the expression of Hexim1 by shRNA (Fig. S4E), and observed a significant reversal of radio-resistance in the irradiated CNE2-IR cells (Fig. S4F). These results showed that inhibition of transcription pausing was an effective way to reverse IR-resistance in cell culture.

**Figure 4.**
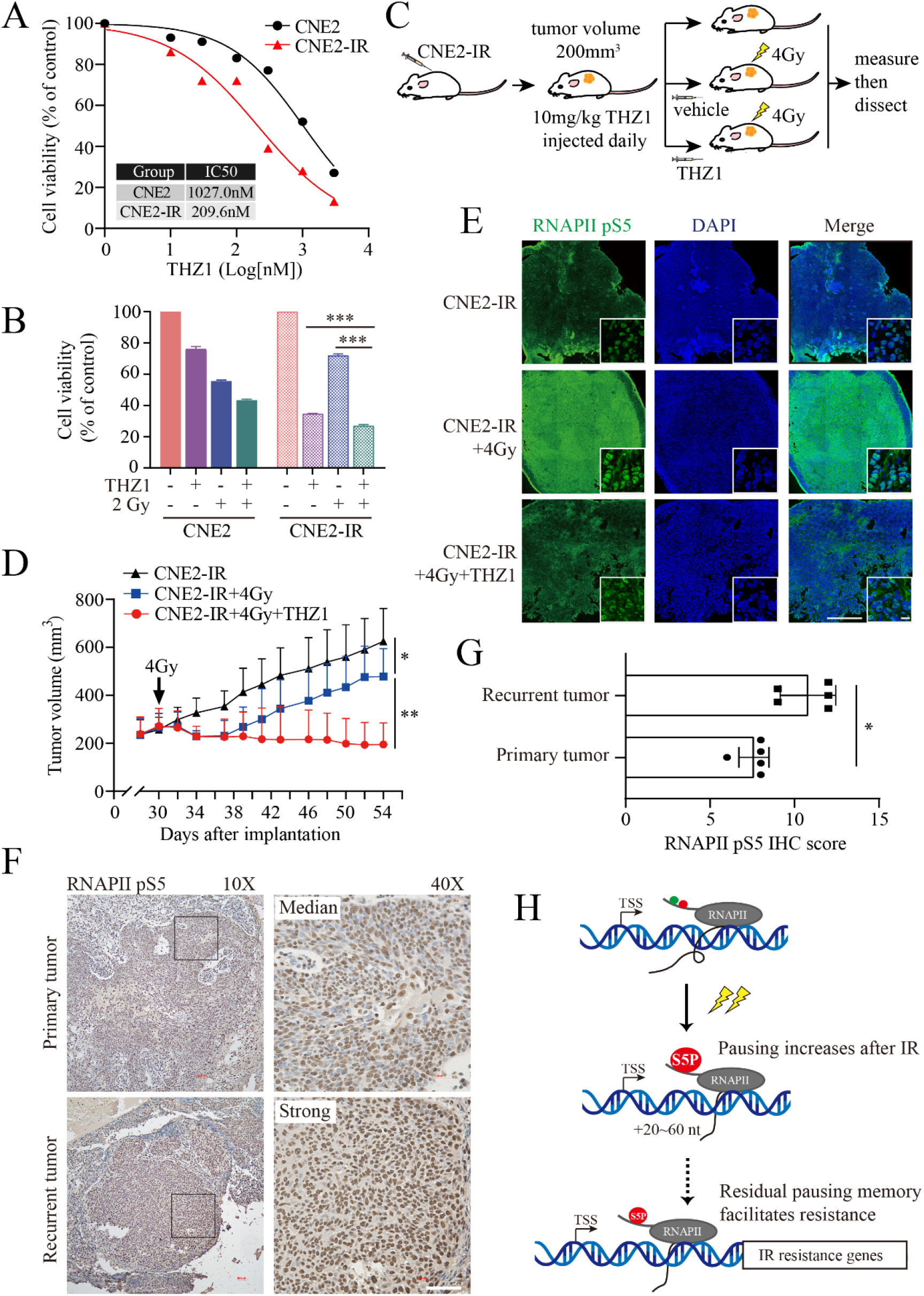
Targeting transcription pausing reverses IR-resistance. (A) Viability assay of CNE2 and CNE2-IR cells treated with different concentrations of THZ1. (B) The combined inhibitory effects of THZ1 and radiation. Data represent mean ± SD. ***p < 0.001 by t-test. (C) Schematic of the experimental procedure testing IR-resistance in xenograft tumor model. (D) Tumor volumes in different experimental groups after radiation. Data represent mean ± SD. n = 5. *p < 0.05, **p < 0.01 by t-test. (E) Immunofluorescence staining of the frozen sections of the mice subcutaneous tumor. RNAPII pS5 is in green, and DAPI in blue. Scale bar: 1000 μm, scale bar in the lower right inset: 10 μm. (F) Examples of immunohistochemical staining for RNAPII pS5 in primary and recurrent nasopharyngeal carcinoma. Scale bar: 50 μm. (G) Semiquantitative comparison of RNAPII pS5 IHC scores between primary and recurrent tumors. Data represent mean ± SD. n = 5. *p < 0.05 by t-test. (H) Working model showing that RNAPII pausing increases after ionizing radiation and the residual pausing memory on a subset of genes contributes to the development of IR-resistance.

To test if inhibition of transcription pausing could be equally effective in vivo, we seeded the radio-resistant CNE2-IR cells subcutaneously into nude mice and generated the xenograft model (Fig. 4C). After the tumor volume reached 200 mm^3^, mice in the experimental groups were injected with vehicle or THZ1 and received 4 Gy of irradiation treatment. The THZ1 was then administrated daily and the tumor volume was measured thrice a week. As shown in Fig. 4D, radiation alone only mildly inhibited the tumor growth, whereas radiation combined with THZ1 treatment completely blocked the tumor growth. We dissected the tumors at the end of the experiment, and prepared frozen sections of the tumor tissues for immunostainings with the RNAPII pS5 antibody (Fig. 4E). As expected, treatment with THZ1 inhibited the radiation-induced elevation of pS5 levels in the tumor tissues (Fig. S4G).

We wanted to know if the observed change in transcription pausing has clinical relevance. We collected 5 paired primary and recurrent nasopharyngeal carcinoma biopsy samples, and performed immunohistochemical analysis to detect the pS5 RNAPII levels (Fig. 4F). By carefully calculating the IHC scores (16), we found the recurrent tumors had a small but steady increase in the pS5 level compared with the primary tumors (Fig. 4G), indicating that the observation made here was of clinical importance.

**Figure S4.**
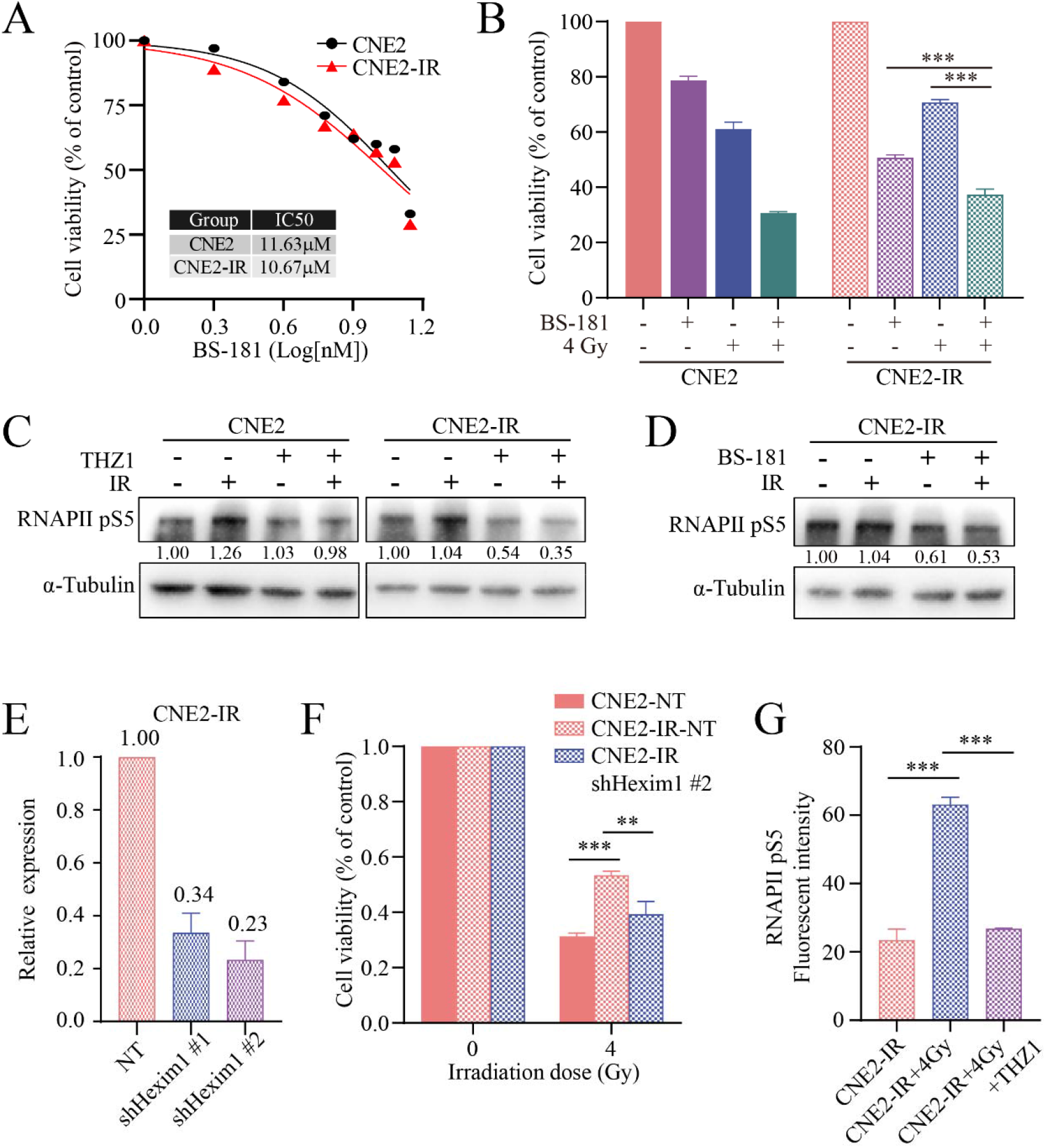
Downregulating transcription pausing improves the effect of radiotherapy. (A) Cell viability assay detecting the sensitivity of CNE2 and CNE2-IR to CDK7 inhibitor BS-181. (B) The combined inhibitory effect of BS-181 with radiation on CNE2 and CNE2-IR cells. Data represent mean ± SD. ***p < 0.001 by t-test. (C-D) Western blot detecting RNAPII pS5 level in CNE2 and CNE2-IR cells treated with radiation (4 Gy) and CDK7 inhibitors. (E) qPCR analyzing Hexim1 mRNA level in CNE2-IR cells transduced with lentiviruses carrying the indicated shRNA. Data represent mean ± SD. (F) The combined inhibitory effect of Hexim1 knockdown with radiotherapy on CNE2 and CNE2-IR cells. Data represent mean ± SD. **p < 0.01, ***p < 0.001 by t-test. (G) Comparison of RNAPII pS5 fluorescent intensity in the indicated experimental groups. ***p < 0.001 by t-test.

## Discussion

In this study, we reported an IR-induced increase of transcription pausing indispensable for the development of radio-resistance (Fig. 4H). The IR exposure evoked an accumulation in pS5 and paused RNAPII. This change of transcriptional kinetics was maintained on a fraction of genes, downregulating their expression and hence facilitating the cells to gain resistance to irradiation. We identified NEK7 as a new resistance gene subjected to such regulation. Knockdown of NEK7 disrupted the bipolar spindle formation, resulting in the prolongation of mitosis and genomic instability which might promote tumor evolution and the development of IR-resistance (3).

The IR-induced increase in pS5 of RNAPII was sensitive to ATM and DNA-PK inhibition, indicating that the observed transcriptional change is an integral part of the cellular responses toward DNA DSBs (17). The relationship between transcription and DSBs seems to be bilateral. On one side, transcriptional elongation at specific loci requires local DNA breaks (23,24), and the release of paused RNAPII promotes cancer-associated translocations, especially around the boundaries of the topologically-associating domains (TADs) (24). On the other side, DNA DSBs induced by restriction enzymes cause a local accumulation of inhibitory H2AK119ub and transcriptional repression in proximity to the DSBs (11,12,24,25). Our finding that IR-induced DSBs elicit a global increase in RNAPII pausing reveals another layer of regulation, adding to our understanding of transcriptional plasticity in DNA damage response.

How DNA DSBs signaling pathways modulate global transcriptional dynamics is the key question awaits future investigation. The answer is likely hidden in the complex post translational modifications of the RNAPII CTD and its regulators. Components in the NELF complex are recruited to DSBs in a RNAPII and PARP-1 dependent manner to repress transcription and promote repair (26). More interestingly Cyclin T, which is required for the activation of CDK9 kinase in P-TEFb, is able to undergo phase separation, forming liquid droplets in vitro that preferentially attract CTD pre-phosphorylated by CDK7 (27). It is tempting to speculate that DNA DSBs and poly(ADP-ribosyl)ation, by regulating the Cyclin T phase separation, modify the transcription cycle and increase RNAPII pausing. The increase in paused RNAPII may reduce transcription elongation and inhibit new initiation of transcription (28).

Lastly, our discovery showed that inhibition of CDK7 reversed the radio-resistance of NPC cells, and that the recurrent NPC patient samples after radiotherapy displayed elevated pS5 of RNAPII, pointing to the possibility of enlisting CDK7 inhibitors as a new class of adjuvant chemo drugs to enhance the efficacy of radiotherapy and thwart treatment resistance. More preclinical and clinical studies are needed to further advance this emerging paradigm.

## Authors’ Contributions

**H. Liu**: Data curation, visualization, investigation, methodology, formal analysis. **C. Yu**: Investigation, methodology, validation. **N. Zhang**: Investigation, resources. **Y. Meng**: Investigation. **C. Huang**: Investigation. **C. Hu**: Resources. **F. Chen**: Investigation, methodology. **Z. Xiao**: Resources. **Z. Zhang**: Resources, supervision. **H. Shao**: Investigation, supervision, funding acquisition. **K. Yuan**: Conceptualization, supervision, funding acquisition, investigation, methodology, visualization, resources, formal analysis, project administration, writing-original draft, writing-review and editing.

## Acknowledgements

We gratefully acknowledge Prof. Liangfang Shen, Prof. Rong Tan, and Dr. Xingming Deng for making reagents and equipment available. We thank Prof. Huasong Lu at Zhejiang University for insightful suggestions, and colleagues in Central South University and members of the Yuan lab for helpful discussions. We apologize for citing reviews instead of original articles due to the limit on the number of references. This project has been supported by the National Natural Science Foundation of China (grants 32170821, 31771589, 91853108 to K.Y., 82104001 to H.S, and 32101034 to F.C), Department of Science & Technology of Hunan Province (grants 2021JJ10054, 2019SK1012, 2018DK2015, 2017RS3013, 2017XK2011 to K.Y, the innovative team program 2019RS1010, 2021JJ41015 to H.S, and 2021JJ41049 to C.Y), Central South University (2018CX032 to K.Y, and the innovation-driven team project 2020CX016). K.Y is supported by the National Thousand Talents Program for Young Outstanding Scientists.

